# When blood needs to flow: The role of the complement system in red blood cell clearance during immune responses

**DOI:** 10.1101/2025.05.23.655770

**Authors:** Clemente F. Arias, Agustín Tortajada, Francisco J. Acosta, Cristina Fernandez-Arias

## Abstract

The complement system is a crucial component of the innate immune response, primarily recognized for its defensive role against pathogens. Increased red blood cell (RBC) mortality due to complement activation during infections is often considered collateral damage of this defensive role. In this study, we propose that this phenomenon results from the active regulation of RBC mortality by the complement system during acute immune responses. Traditionally, RBC homeostasis has been explained by the need to ensure adequate oxygen delivery. However, RBCs are also critical determinants of blood viscosity, which directly influences tissue perfusion. Efficient immune responses require increased blood flow to infected tissues, but changes in plasma proteins during acute-phase responses can increase blood viscosity, hindering perfusion. We suggest that the complement system selectively removes less elastic RBCs, compensating for the rise in plasma viscosity and optimizing tissue perfusion during infections. Our findings reveal a previously unrecognized role of the complement system in regulating blood flow, highlighting the intricate balance between complement’s homeostatic and defensive functions in maintaining physiological stability.

## Introduction

The complement system is a critical component of immunity, enhancing the clearance of pathogens, driving inflammation, and directly lysing foreign cells [1]. This system functions as a highly coordinated network of proteins and glycoproteins that detect and respond to invading agents. Different complement pathways converge in the formation of C3 convertases [2], enzymatic protein complexes, with serine protease activity, that cleave the C3 component into C3a and C3b fragments [3]. This cleavage exposes a highly reactive thioester group in C3b, allowing it to covalently bind to nearby surfaces, such as pathogen or host cell membranes [4, 5].

The binding of C3b to a pathogen’s membrane has two possible outcomes. First, it can trigger a cascade of reactions that forms membrane attack complexes (MACs), non-enzymatic structures that perforate the cell membrane, lysing and killing the cell [6, 7]. Additionally, C3b is a potent opsonin that tags cells for phagocytosis by macrophages and Kupffer cells [2].

The majority of complement proteins, including C3, are continuously synthesized by hepatocytes in the liver and released into the bloodstream, ensuring that the complement system remains ready to rapidly detect and respond to infections [8, 9]. However, this constant presence of complement proteins in circulation also means that RBCs are continuously exposed to them, potentially leading to pathological interactions. This risk is particularly evident in RBCs lacking complement regulators such as CD55 and CD59—membrane proteins that inhibit different steps of complement activation [10, 11]. The absence or dysfunction of these regulators on the cell membrane can lead to the accumulation of C3b and the subsequent formation of MACs, ultimately lysing RBCs in the bloodstream, a condition known as intravascular hemolysis [12, 13].

In contrast, healthy RBCs with functional complement regulators are typically spared by complement proteins. Instead, they are phagocytosed in the liver and spleen once they reach a critical age—approximately 120 days in humans and 60 days in mice. As they pass through the hepatic and splenic sinusoids, RBCs come into close contact with phagocytes [14]. Macrophage receptors recognize specific molecules on the RBC membrane, such as phosphatidylserine (PS) and CD47, which act as pro- and anti-phagocytic signals, respectively [15]. The selective removal of aged RBCs is determined by the dynamic balance of these signals on the RBC membrane [14]. In young RBCs, high levels of CD47 expression inhibit phagocytosis. As RBCs age, they gradually lose CD47 and accumulate pro-phagocytic signals, which ultimately marks RBCs for destruction in the liver or spleen [16–18].

It is commonly assumed that complement proteins overlap with other mechanisms that enhance phagocytosis, promoting the clearance of RBCs from circulation. For example, complement proteins can bind to RBCs opsonized with autoantibodies, facilitating their recognition and removal by phagocytes in the liver [17]. From this perspective, the complement system would function as a failsafe mechanism, ensuring the clearance of senescent or dysfunctional RBCs from the bloodstream [19].

In this work, we propose that the role of the complement system extends beyond the elimination of aged or damaged cells. Specifically, we suggest that it plays a fundamental role in RBC lifespan by acting in conjunction with PS and CD47 to regulate the physiological removal of healthy RBCs. From this perspective, the complement system is integrated into the normal turnover of RBCs, with continuous C3 deposition and generation of different opsonin fragments, which are recognized through different receptors on phagocytes in distinct tissue locations. This deposition continuously occurs throughout their life cycle, and not only when they are marked for phagocytosis.

Using mathematical modeling, we explore the implications of this continuous opsonization and demonstrate its essential role in RBC homeostasis, particularly during acute responses to infections. We propose that this mechanism actively shortens RBC lifespan during inflammation, thereby reducing the overall number of circulating cells. This reduction would play a key role in regulating tissue perfusion during immune responses.

## Results

### The dynamics of complement protein on red blood cell membranes

C3 convertases can be produced via three different pathways. In the classical pathway, they are formed when the C1q protein binds to IgG- or IgM-containing immune complexes [20]. In the lectin pathway, this process is triggered by lectins like the mannose-binding lectin (MBL), which recognizes and binds carbohydrate or glycoprotein components commonly found on the surface of pathogenic microorganisms [21]. Finally, the alternative pathway keeps a fluid-phase C3 convertase basal level, due to the spontaneous hydrolysis of C3 into C3(H20), in a process called “tick-over”. Fluid-phase C3 convertase maintains a low level formation of C3b, which is rapidly inactivated by complement regulators [22, 23].

The lectin pathway primarily targets foreign cells and has minimal impact on RBCs. In contrast, the alternative pathway, due to the constitutive activation, targets both pathogens and host cells [24], leading to a continuous formation of C3b on normal RBCs. Additionally, natural antibodies that recognize RBC self-antigens circulate in the blood [25, 26], activating the classical pathway and further promoting C3b deposition on RBCs [27].

Complement inhibitors such as CD55 and CD59 prevent C3 convertase formation and the MAC formation on RBCs, favoring instead the conversion of C3b to iC3b and its subsequent cleavage to C3dg by serin protease factor I and cofactors factor H and CR1 (Fig. 1.A). The effects of RBC opsonization with iC3b and C3dg remain poorly understood. Complement receptors, such as CR-Ig, CR3, and CR4 can recognize both opsonins, suggesting that they might trigger similar cellular responses [28–33]. However, these receptors preferentially recognize iC3b rather than C3dg [32]. Consequently, iC3b-opsonized RBCs are more likely to be phagocytosed by Kuppfer cells than those opsonized with C3dg [17, 34]. C3dg-coated RBCs can still be phagocytosed under particular circumstances, particularly by activated monocytes [35, 36].

**Figure 1.**
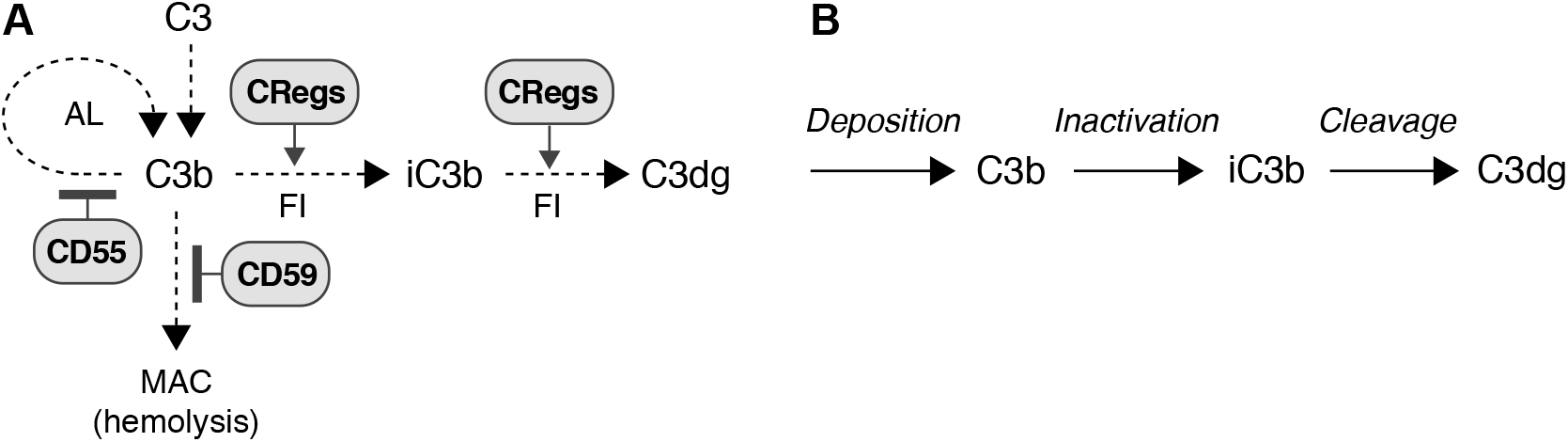
Dynamics of complement proteins on RBC membranes. A) The binding of C3b to the RBC surface can lead to three possible outcomes. First, it may trigger an amplification loop (AL), further increasing C3b deposition. Second, C3b can contribute to the formation of C5 convertases, which cleave C5 into C5a and C5b, initiating the membrane attack complex (MAC) and leading to RBC lysis. Third, C3b can be inactivated by Factor I, with the help of the soluble regulator factor H and membrane cofactors such as CR1 in humans or Crry in mice, forming iC3b. Factor I can further cleave iC3b using CR1 as cofactor to produce C3dg, which remains bound to the RBC membrane. On healthy RBCs, the amplification loop and MAC formation are inhibited by complement regulators CD55 and CD59, respectively. B) This work focuses on C3b deposition, its inactivation to iC3b, and subsequent cleavage to C3dg, as these are the main opsonins impacting RBC phagocytosis.

In this work, we hypothesize that iC3b and C3dg convey distinct signals to phagocytes, regulating alternative phagocytosis pathways with different roles under physiological and pathological conditions. Specifically, we propose that C3dg mediates the phagocytosis of RBCs in cases of blood extravasation resulting from tissue injury. The rapid clearance of extravasated RBCs is essential to prevent oxidative damage caused by hemoglobin release, which can disrupt the local tissue environment and exacerbate inflammation [37]. We suggest that activated monocytes rely on C3dg signals to perform this task. These monocytes are promptly recruited to sites of infection or injury [38].

Healthy RBCs only interact with activated monocytes in case of extravasation. Therefore, the C3dg is inactive in the physiological context of RBCs within the circulatory system. We propose that, in this context, iC3b acts as a homeostatic pro-phagocytic signal, acting in conjunction with PS and CD47 in the identification and selective removal of RBCs during homeostatic turnover.

To explore the implications of this hypothesis, we will next model the dynamics of complement proteins on the membrane of individual RBCs. Specifically, we will focus on the deposition of C3b on healthy RBC membranes, its inactivation to iC3b, and its subsequent cleavage to C3dg (Fig. 1.B).

### A model of complement dynamics on RBC membranes

The dynamics of complement proteins on the membrane of RBCs, as outlined in Fig. 1.B, can be modeled by the following equations:

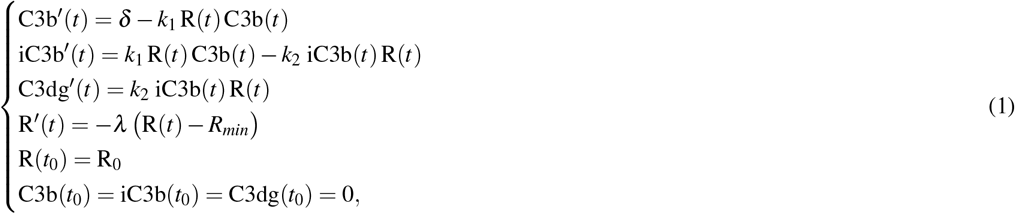

where *d*(*t*) represents C3b deposition on the cell membrane at time *t*, and *k*_1_ and *k*_2_ are the rate constant for the conversion from C3b to iC3b, and the rate constant for the cleavage of iC3b to C3dg, respectively. R denotes the expression of complement regulators in the RBC membrane, such as CR1 and Crry, that are essential for the inactivation of C3b and the cleavage of iC3b by Factor I. For simplicity, we assume that the plasma concentration of Factor I and factor H remain constant during the life span of RBCs. The change of complement regulator levels over time reproduces the observed dynamics of CR1 in the membrane of aging RBCs, which follow an exponential decay model with offset [39]. Parameters *λ* and *R*_*min*_ determine the shape of this decay (Fig. 2.A), and R_0_ denotes the level of complement regulators in the membrane of the RBC at the moment of its formation (labelled as *t*_0_).

**Figure 2.**
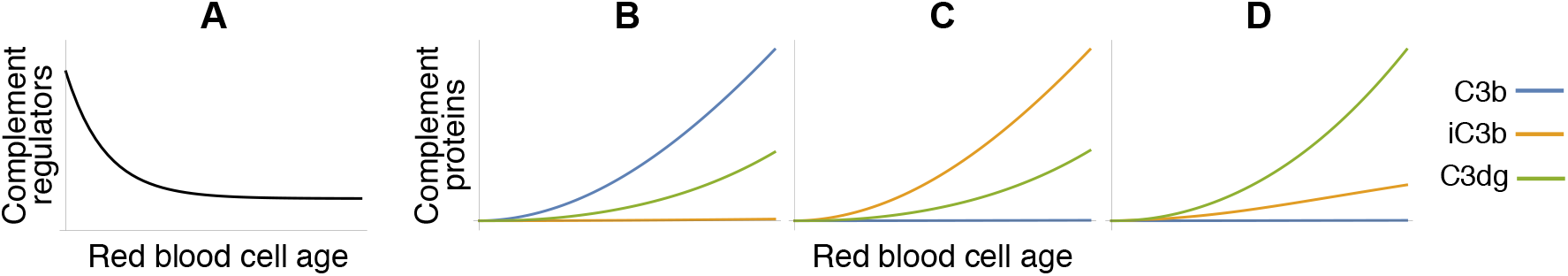
Dynamics of model 1. A) The model assumes that complement receptor expression on the RBC membrane declines exponentially with cell age, reaching a minimum value represented by the parameter *R*_*min*_ [39]. B-D) Numerical simulations of the model under different parameter values. B) When the rate of C3b deposition exceeds the rate of inactivation (*δ > k*_1_), C3b accumulates on the RBC membrane. C, D) Under conditions where *k*_1_ *> δ* and *k*_1_ *> k*_2_, smaller *k*_2_ values result in iC3b predominance (C), while larger *k*_2_ values favor C3dg prevalence (D). In both cases, C3b accumulation is negligible.

We next use equations 1 to analyze the qualitative dynamics of complement activation and regulation on RBC membranes, as well as their impact on RBC lifespan. The exact parameter values are not critical to this analysis. The model’s simplicity allows for a straightforward exploration of its behavior and the derivation of robust conclusions that are independent of specific numerical values.

We begin by defining the function *d*(*t*), which represents the binding of C3b to the cell membrane at any given time *t*. C3b deposition increases with RBC age. This process is facilitated by natural antibodies recognizing self-antigens on the RBC membrane, a phenomenon that intensifies in older cells due to higher antibody accumulation on their surfaces [40]. Additionally, young RBCs are coated with sialic acid, which are recognized by Factor H, a major plasma complement regulator [41, 42]. Factor H inhibits the alternative complement pathway’s amplification loop (Fig. 1.A), preventing local C3b production [43, 44]. As the RBC ages, its sialic acid coating diminishes [45], progressively increasing C3b deposition. For simplicity, the age-dependent increase in C3b deposition is modeled as:

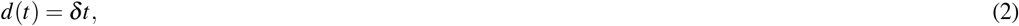

where *δ* is a positive parameter that condenses the combined contributions of the different complement pathways to C3b deposition. We assume that the value of this parameter is greater for higher plasma concentrations of C3 and C3(H2O), reflecting the dependency of C3b deposition on systemic complement availability.

Replacing *d*(*t*) with this expression in equations 1 describes the dynamics of C3b, iC3b, and C3dg on the membrane of individual RBCs throughout their lifespan. Numerical simulations of this model for different parameter values yield alternative outcomes in terms of the relative abundance of each complement molecule over time (Figs. 2.B-D).

Scenarios where the rate of C3b deposition is greater than the rate of inactivation to iC3b result in the accumulation of C3b (Fig. 2.B), which, in living RBCs, would lead to the formation of MACs and subsequent cell lysis [46, 47]. Under physiological conditions, RBCs are primarily removed through phagocytosis in the liver and spleen, and lysis occurs only sporadically [17]. This indicates that C3b inactivation must outpace its deposition to prevent its accumulation on the cell membrane.

On the other hand, empirical evidence indicates that C3b inactivation occurs faster than iC3b cleavage [48–50], which, in the context of model 1, implies that *k*_2_ *< k*_1_. Under this constraint, smaller values of *k*_2_ result in the predominance of iC3b (Fig. 2.C), whereas larger values favor the prevalence of C3dg (Fig. 2.D).

In the remainder of this text, we will use the complement dynamics shown in Fig. 2.D, which prevent the excessive accumulation of C3b and reproduce the progressive opsonization of RBCs with iC3b and C3dg. Our focus will be on iC3b and its interactions with PS and CD47, so the abundance of C3dg does not affect the conclusions drawn from model 1. In the following sections, we will use this model to explore the impact of the complement system on RBC lifespan.

### The role of complement in RBC lifespan

In previous works, we introduced a formal model of RBC lifespan determination based on the dynamics of CD47 and PS on the RBC membrane (see [51] and [52] for details). Denoting the levels of CD47 and PS by *C*(*t*) and *P*(*t*), respectively, this model is given by:

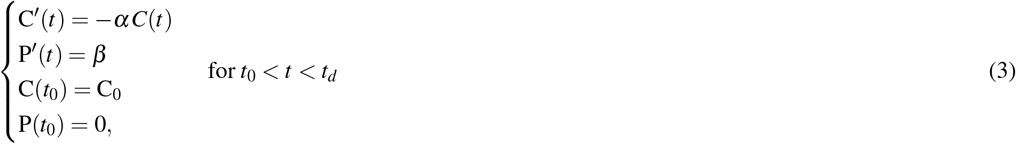

where *α* and *β* are positive parameters, and *t*_0_ and *t*_*d*_ represent the moments of the RBC formation and death respectively.

For simplicity, pro-phagocytic signal molecules, such as natural autoantibodies that target different antigens of RBCs, promoting their phagocytosis, are not explicitly included in the model. However, the dynamics of these signals are analogous to PS, increasing with RBC age [25]. Consequently, the PS equation in the model can be interpreted as a representative proxy for generic pro-phagocytic molecules.

In this work, we generalize model 3 as follows:

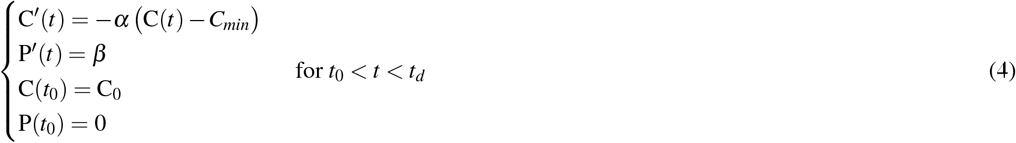

This model assumes that the dynamics of CD47 are analogous to those observed for CR1 on the membrane of human RBCs. The progressive loss of complement regulators is largely caused by vesiculation [53, 54], a process by which portions of the membrane are shed as vesicles [55, 56]. As the RBC ages, CR1 declines exponentially until eventually stabilizing at a minimum value [39] (Fig. 2.A). Model 4 assumes a similar pattern of decline, with *C*_0_ and *C*_*min*_ representing the initial and final levels of CD47 expression, respectively. We remark that models 3 and 4 coincide if *C*_*min*_ = 0.

To model the effect of complement on RBC lifespan, equations 1 and 4 can be integrated as follows:

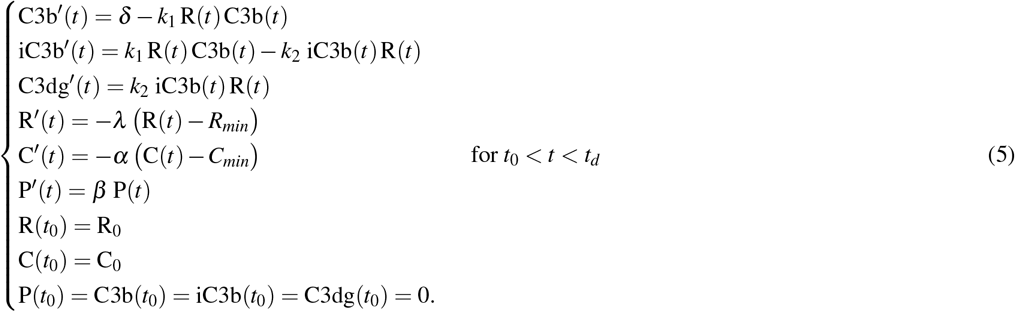

This model captures the simultaneous dynamics of CD47, PS, and complement proteins on the membrane of an individual RBC, from its formation at time *t*_0_ to its eventual death at time *t*_*d*_.

These molecules interact with specific macrophage receptors. PS is recognized by Tim-1, Tim-4, and Stabilin-2, among others [18, 57, 58], whereas CD47 primarily interacts with SIRP-*α* [59]. On the other hand, iC3b is recognized by complement receptors highly expressed by macrophages, such as CR3, CR4, and CR-Ig [30, 47, 60]. The interaction between RBC signals and membrane receptors determines the information perceived by macrophages during their contact with RBCs.

In this work, we hypothesize that iC3b is an additional pro-phagocytic signal, acting in conjunction with PS and CD47 to determine whether RBCs should be phagocytosed as they pass through the hepatic and splenic sinusoids. This implies that, in their interactions with RBCs, macrophages evaluate the difference between pro-phagocytic stimuli conveyed by PS and iC3b, and anti-phagocytic stimuli conveyed by CD47. If this difference exceeds a given threshold, the RBC is phagocytosed. Otherwise, it continues to circulate in the bloodstream.

This condition can be expressed as follows:

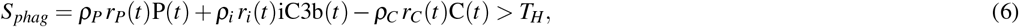

where *T*_*H*_ is the signal threshold that triggers phagocytosis, and *S*_*phag*_ represents the phagocytic signals perceived by the macrophage in its interaction with the RBC. The number of CD47, PS, and iC3b receptors are denoted by *r*_*C*_, *r*_*P*_, and *r*_*i*_, respectively. Parameters *ρ*_*CD*47_, *ρ*_*PS*_, and *ρ*_*CR*_ quantify how the molecules on the RBC surface influence the macrophage’s response. We will refer to this phagocytosis pathway as homeostatic phagocytosis.

Red pulp macrophages also detect and phagocytize RBCs with low CD47 levels, regardless of their prophagocytic levels [61, 62]. This mechanism triggers autoimmune responses against circulating RBCs, which helps explain the normal presence of natural autoantibodies targeting RBC antigens in the blood [52, 62]. The trigger of this phagocytosis pathway, which we labeled immune phagocytosis in a previous work [51], can be expressed as:

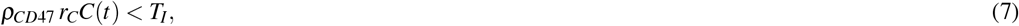

where *T*_*I*_ denotes the lower boundary of CD47 that triggers immune phagocytosis of the RBC.

The comparison between models 4 and 5 shows that the mechanism of RBC lifespan determination based on CD47 and PS does not qualitatively change with the inclusion of iC3b as an additional pro-phagocytic signal (Fig. 3:A). In both models, varying levels of CD47 expression in newly formed cells influence not only the lifespan of the RBC but also its phagocytosis pathway (homeostatic or immune) (Figs. 3.B,C).

**Figure 3.**
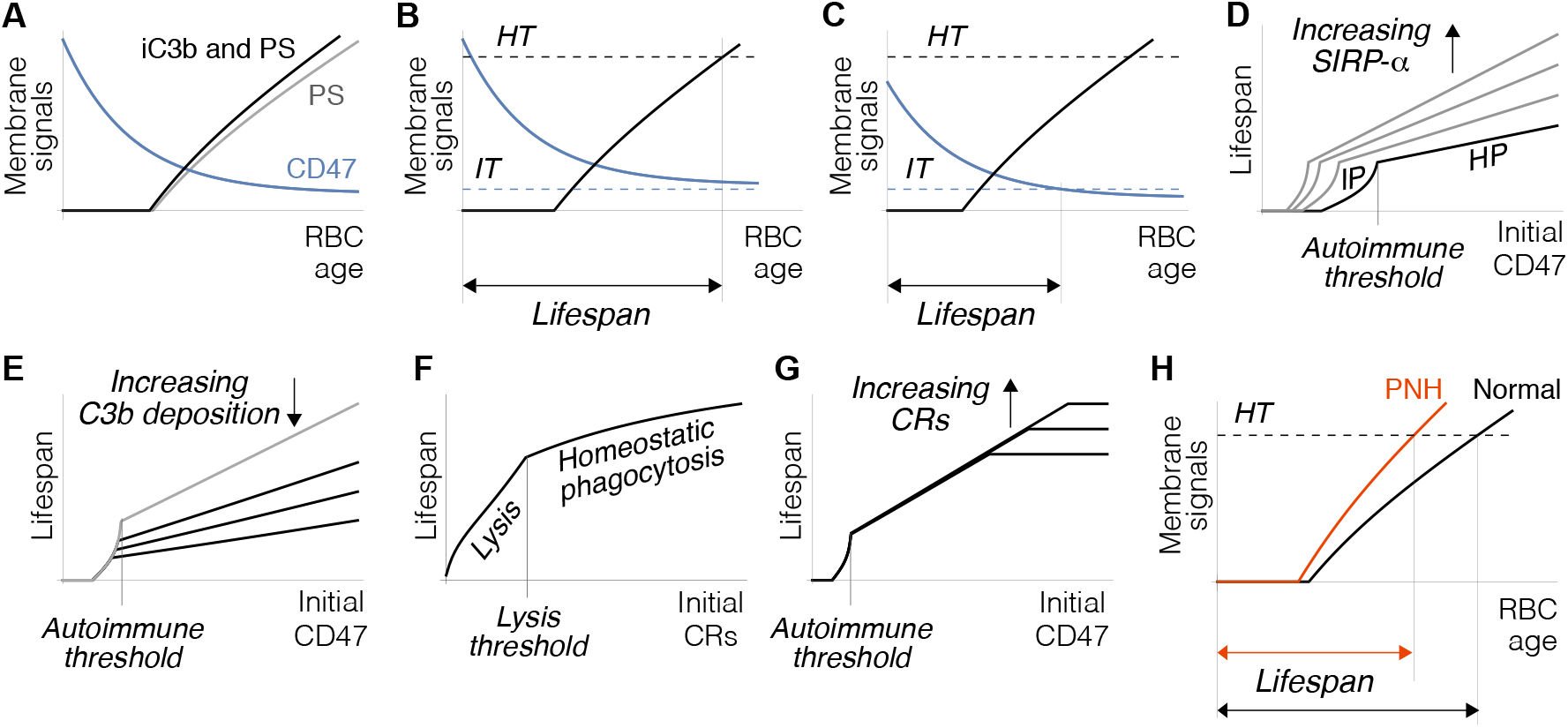
Dynamics of model 5. A) Dynamics of CD47 and phagocytic signals (PS - CD47, gray; PS + iC3b - CD47, black). Incorporating iC3b in the model does not qualitatively alter the dynamics of phagocytic signals delivered by RBCs to macrophages. B, C) The dynamics of membrane signals determine the lifespan and phagocytosis pathway of RBCs. Phagocytosis occurs when phagocytic signals exceed a certain threshold (*TH*) (B) or when CD47 levels fall below a critical value (*IT*) (C). Under the second condition, RBC phagocytosis may trigger an autoimmune response against circulating RBCs. D) The lifespan and phagocytosis pathway of RBCs are determined by the initial levels of CD47 expression at the time of the cell formation. Higher CD47 levels lead to homeostatic phagocytosis (HP), whereas lower levels trigger immune phagocytosis (IP). The CD47 value at which this transition occurs is labelled as the autoimmune threshold, and is modulated by the expression levels of SIRP-*α* receptors on macrophages (gray lines). E) Lifespan and the autoimmune threshold can be further modulated by C3b deposition levels. The gray line represents a scenario without C3b deposition. F) The levels of complement regulators at the time of cell formation (denoted by *R*_0_ in the model) affect the fate and lifespan of RBCs. Cells with low levels of complement regulators undergo lysis, whereas those with high levels are phagocytized by macrophages. We define the lysis threshold as the value of regulators that determines the transition between these two removal pathways. G) Complement proteins establish an upper limit to RBC lifespan. This limit is higher for RBCs formed with greater levels of complement regulators on their membranes. H) Model 5 predicts hemolysis in PNH patients treated with eculizumab. Increased C3b deposition accelerates the homeostatic phagocytosis pathway (red), leading to a shortened RBC lifespan compared to normal conditions (black). *HT* : homeostatic threshold.

RBCs with higher levels of CD47 at the time of their formation undergo homeostatic phagocytosis at progressively shorter lifespans (Fig. 3.D). In contrast, RBCs with lower CD47 levels are removed via the immune pathway instead (Fig. 3.D). The critical value of CD47 expression that triggers this transition is modulated by SIRP-alpha expression in macrophages [52] (Fig. 3.D). Although this effect of CD47 can be further modulated by complement proteins (Fig. 3.E), it exists independently of the complement system (see [51, 52]).

Therefore, including complement proteins in model 4 appears redundant, as they do not significantly alter the overall dynamics of RBC lifespan regulation. However, their inclusion introduces important differences. For instance, according to model 5, the complement system establishes an upper boundary for RBC lifespan. As noted earlier, the accumulation of C3b on the RBC membrane results in the formation of MACs, which ultimately leads to cell lysis. The trigger of lysis by the accumulation of C3b beyond a critical threshold *T*_*L*_ can be expressed as:

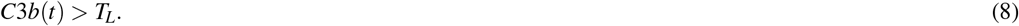

If this condition is included in model 5, RBC lifespan is ultimately constrained by the accumulation of C3b. This constraint depends on the number of complement regulators present on the RBC membrane at the time of its formation. In RBCs formed with low levels of complement regulators, increased C3b deposition may lead to the lysis of the cell in the bloodstream. Conversely, RBCs formed with higher levels of complement regulators are protected from complement-mediated cell lysis and are removed via the homeostatic phagocytosis pathway (Fig. 4.F). We define the lysis threshold as the critical level of complement regulators that determines whether the RBC will undergo lysis (Fig. 4.F).

**Figure 4.**
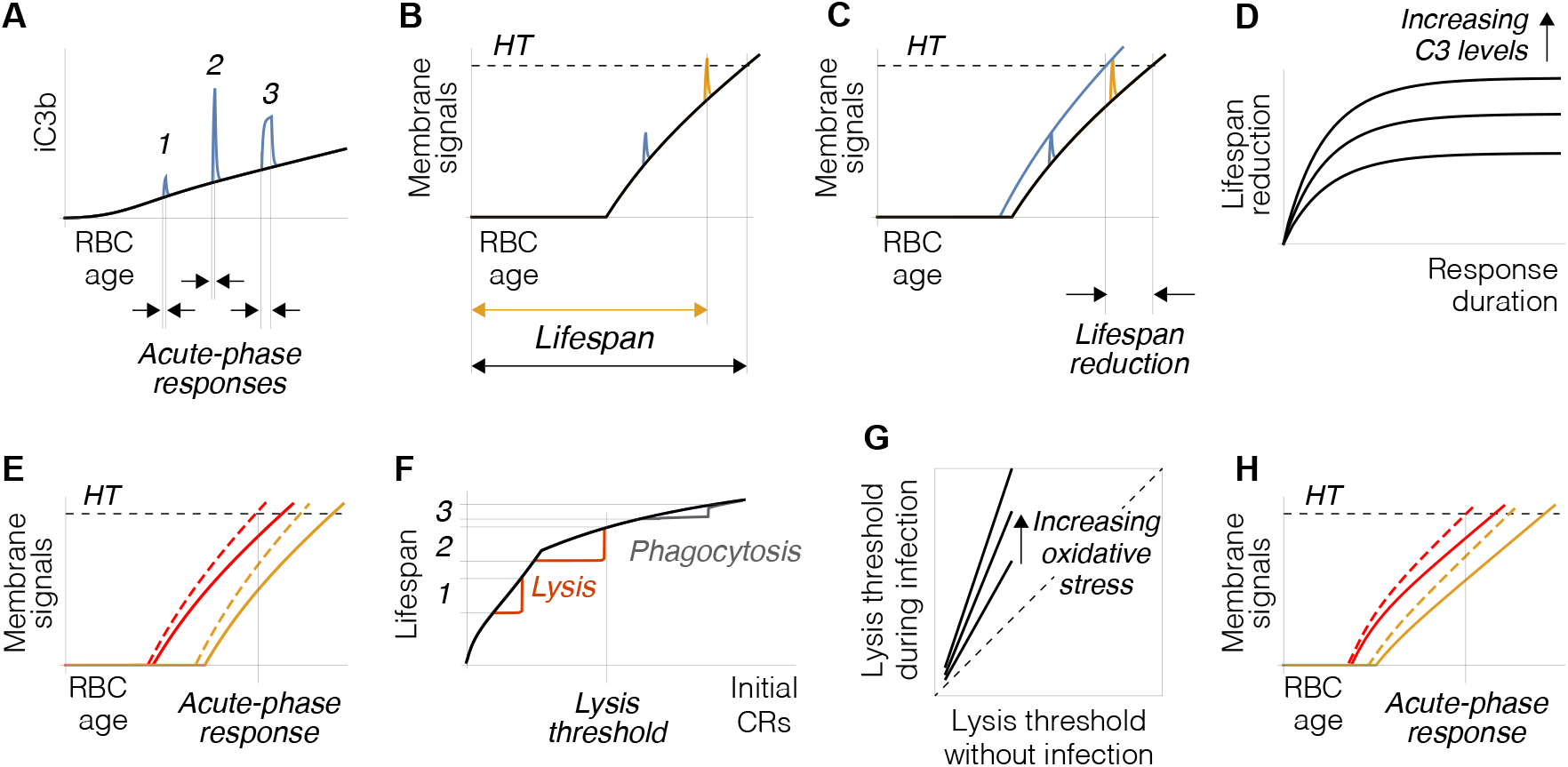
Dynamics of model 5 during acute-phase responses. A) Acute-phase responses cause a transient increase in iC3b, which amplifies the phagocytic signals delivered by the RBCs to macrophages. The peak of this increase depends on the timing, duration, and intensity of the response (gray lines). Once the response concludes, phagocytic signals return to baseline levels (black line). B) The effect of the response critically depends on the age of the RBC. In older cells (blue line), increased phagocytic signals may trigger phagocytosis, leading to premature cell death (*HT* : homeostatic threshold). C) Acute responses cause a reduction in RBC lifespan, primarily affecting older cells. The blue line interpolates the peaks of phagocytic signals for identical responses at different RBC ages. D) The reduction in lifespan increases with higher C3 plasma levels and longer durations of the response. E) RBC age is not the only determinant of premature death. The red and orange lines represent the dynamics of phagocytic signals in two RBCs formed at the same time, with lower (red) and higher (orange) levels of CD47. An acute response may trigger premature death in one but not in the other. Dashed colored lines interpolate the peaks of phagocytic signals at different times. F) Acute responses accelerate the lysis of RBCs with low initial levels of complement regulators (1). They may also alter the lysis threshold, causing RBCs that would typically be phagocytosed in the absence of infection to undergo lysis instead (2). RBCs with higher levels of complement regulators undergo premature death through phagocytosis (3). G) Changes in the lysis threshold caused by acute-phase responses increase with oxidative stress. Accelerated loss of complement regulators due to oxidative stress makes lysis more likely. H) Increased oxidative stress can also accelerate homeostatic phagocytosis. Red and orange lines represent the effect of higher (red) and lower (orange) oxidative stress on the dynamics of phagocytic signals in two RBCs formed at the same time. Increased oxidative stress makes premature death more likely. Dashed colored lines interpolate the peaks of phagocytic signals at different times.

Apparently, this constraint represents a critical limitation on the effect of CD47 on lifespan—an increase in CD47 would not extend lifespan but rather heighten the risk of lysis (Fig. 3.G). However, empirical evidence suggests that EPO enhances the expression of both CD47 [52] and CR1 [63–65] on newly formed RBCs. According to model 5, this implies that RBCs predestined to live longer are also better protected from C3b-mediated lysis than those with shorter expected lifespans (Fig. 3.G).

Model 5 also accounts for some of the consequences of complement dysregulation on RBC lifespan. Deficiencies in complement inhibitors CD55 and CD59 dramatically increase C3b deposition, promoting MAC formation and leading to intravascular hemolysis [66]. This condition, known as paroxysmal nocturnal hemoglobinuria (PNH), is characterized by abnormally accelerated RBC destruction [66, 67].

Eculizumab, a monoclonal antibody that inhibits terminal complement activation, significantly reduces hemolysis in PNH patients by preventing the formation of C5b and inhibiting MAC assembly [66, 68]. However, eculizumab does not prevent the complement amplification loop (see Fig. 1.A), and the subsequent increase in C3b deposition [69]. In these circumstances, model 5 predicts a reduction in RBC lifespan, consistent with the intravascular hemolysis observed in eculizumab-treated PNH [66, 70] (Fig. 3.H). In summary, this section integrates the model of successive C3b inactivation and cleavage on the RBC membrane with the mechanism of RBC lifespan determination. We have shown that including iC3b as a pro-phagocytic signal in the model does not qualitatively differ from macrophage phagocytosis decisions based solely on CD47 and PS. However, it explains the onset of intravascular hemolysis observed in PNH as a consequence of increased C3b deposition. Moreover, it suggests that RBC lifespan is constrained by an upper limit that varies depending on the levels of C3 and complement regulators.

### The role of complement in RBC lifespan during acute phase responses

C3 is an acute-phase protein, meaning that its concentration increases by at least 50% during inflammation and immune responses to pathogens [71, 72]. This boost in C3 levels promotes C3b deposition, leading to a peak of iC3b on circulating RBCs (Fig. 4.A). As noted earlier, the rate of C3b inactivation is greater than the rate of iC3b cleavage to C3dg. Consequently, the excess C3b is rapidly converted into iC3b [73], which is then more slowly cleaved into C3dg [48, 49]. This results in the transient accumulation of iC3b. Once the acute-phase response concludes, C3 levels return to normal pre-infection levels in the blood, and excess iC3b is eventually cleaved to C3dg (Fig. 4.A). According to model 5, the peak of iC3b has no significant effect in young RBCs. In contrast, increased iC3b levels may lead to a premature death in older cells (Figs. 4.B,C), an effect that depends on both the intensity and duration of the response (Fig. 4.D). The risk of premature death is greater in RBCs with low initial C47 levels, which have shorter predefined lifespans (Fig. 3.C). As a result, they are more likely to be removed, while RBCs of the same age with higher CD47 levels can survive (Fig. 4.E).

The buildup of C3b on the RBC membrane during acute-phase responses also increases the risk of intravascular lysis. Model 5 suggests that elevated C3 plasma levels accelerate the lysis of RBCs with low levels of complement regulators (Fig. 4.F) and raise the lysis threshold, causing RBCs that would normally be phagocytosed to undergo premature lysis instead (Fig. 4.F).

Remarkably, the organism anticipates the increase in hemolysis caused by the complement system during immune responses. The rupture of RBCs in the bloodstream releases hemoglobin, which can lead to oxidative stress and tissue damage [12]. This is counteracted by haptoglobin, which binds to free hemoglobin released during hemolysis, and ceruloplasmin, which binds to free iron in the blood [12, 74, 75]. The levels of these proteins significantly increase during acute-phase responses, helping to minimize the detrimental consequences of increased hemolysis [71, 74, 76]. Model 5 also predicts an increase in the susceptibility to premature death with oxidative stress, which exacerbates vesiculation [77], amplifying the loss of complement regulators from the cell membrane. This can accelerate lysis (Fig. 4.G), and trigger complement-mediated phagocytosis at earlier stages of the RBC life cycle (Fig. 4.H).

The conclusion drawn from these results is that increased C3 levels during acute-phase responses lead to a reduction in the number of RBCs in the blood. Given that approximately 1% of circulating RBCs are removed daily during normal turnover, a reduction in lifespan by even a few days can result in a significant decrease in RBC population size. The increase in RBC mortality during immune responses and inflammation is well documented [78–81]. RBCs killed by the complement system during infections are often regarded as bystander victims of an immune response primarily targeting pathogens, rather than RBCs themselves [79].

Model 5 provides a novel and compelling perspective on this observation. It suggests that the death of RBCs by complement proteins is not random, but rather the result of a highly regulated process. Counterintuitive as it may seem, this appears to be a purposeful response, not merely a collateral effect. What could be the physiological rationale behind this behavior?

We propose that this is an active mechanism to enhance tissue perfusion during infections. The early stages of acute-phase responses have two opposing effects on blood flow to infected tissues. On one hand, inflammation typically leads to the release of various signaling molecules (e.g., histamine or bradykinin) that dilate blood vessels and increase permeability, thereby enhancing perfusion in affected areas [82]. Fever may further contribute to enhanced perfusion, as blood viscosity decreases with temperature [83, 84]. On the other hand, immune responses significantly increase the plasma levels of proteins such as globulins and fibrinogen, which can raise blood viscosity, making it more resistant to flow and hindering perfusion [74].

As a non-Newtonian fluid, blood’s viscosity varies with shear rate [85, 86], which depends on the velocity of flow and the diameter of blood vessels. Consequently, the factors influencing blood viscosity differ across various regions of the circulatory system [85–87]. In small capillaries, the primary site of tissue perfusion, blood viscosity depends on plasma viscosity, hematocrit, and the mechanical properties of RBCs [83, 85, 88]. Therefore, changes in plasma protein content during acute-phase responses, which can increase plasma viscosity by up to 20% [89], may critically compromise perfusion when it needs to be enhanced.

We propose that the complement system helps offset this effect, contributing to the reduction of blood viscosity. Removing RBCs from the bloodstream is an effective way to rapidly lower hematocrit, a major determinant of blood viscosity [90]. Additionally, complement targets RBCs approaching the end of their lifespan. Aging RBCs may lose up to 10% of their membrane through vesiculation, leading to increased stiffness [53, 56, 91]. Less elastic RBCs may have difficulty deforming to pass through narrow vessels, impairing blood flow and locally raising blood viscosity [92]. We suggest that by selectively removing these cells, the average elasticity of the RBCs in circulation increases, further contributing to lower blood viscosity and improved perfusion.

## Discussion

In this work, we present a theoretical model that explores the role of the complement system in regulating RBC lifespan, offering insights into its potential significance in both physiological and pathological contexts. We propose that different complement proteins convey distinct signals to the phagocytes responsible for RBC clearance. Specifically, we suggest that C3dg tags extravasated RBCs for rapid removal in injured tissues, while iC3b, together with CD47 and PS, facilitates the routine turnover of healthy RBCs under physiological conditions. Incorporating iC3b as an additional pro-phagocytic signal does not significantly alter the homeostatic removal of senescent RBCs. The redundancy of iC3b’s role predicted by our model is confirmed by the observation that both RBC lifespan and phagocytosis remain normal in mice deficient in C3 protein [40]. This does not imply that complement proteins are irrelevant or merely accessory in the phagocytosis of RBCs. Rather, our model suggests that they play a fundamental role in regulating RBC homeostasis, particularly during infections.

The deposition of C3b on the membrane of healthy RBCs is not only influenced by intrinsic cellular factors, such as the rate of vesiculation or the loss of complement regulators. It crucially depends on plasma levels of complement proteins. Under normal circumstances, blood C3 levels remain relatively constant. However, as an acute-phase protein, its concentration sharply increases during infections or inflammation [71, 72], resulting in heightened C3b deposition and subsequent accumulations of iC3b on RBC membranes. According to our model, this can cause the premature phagocytosis of RBCs approaching the end of their lifespan.

The transitory increase in plasma C3 levels has additional effects. First, although C3b primarily targets pathogens for opsonization, it can also opsonize RBCs. However, the contribution of C3b to phagocytosis is negligible compared to that of iC3b (see Fig. 1.D). More importantly, elevated C3b levels may promote RBC lysis in the bloodstream.

Therefore, our model predicts the observed increase in RBC mortality observed during infections [78]. This is the consequence of enhanced phagocytosis and lysis triggered by acute immune responses, primarily affecting RBCs close to the end of their lifespan. These cells are likely to be stiffer due to the progressive loss of cell membrane through vesiculation [19]. However, it is important to emphasize that RBCs targeted by increased C3 levels during acute-phase responses are fully functional cells that would otherwise survive longer in the absence of infection, not necessarily defective or dysfunctional (see Fig. 3.B).

In this work, we propose that by selectively removing stiffer RBCs from the bloodstream, the complement system simultaneously increases the average elasticity of RBCs in circulation and reduces hematocrit during acute-phase responses. Such a mechanism would necessarily lower blood viscosity in the microcirculation, thereby improving blood flow to infected tissues.

This approach represents a significant shift from the prevailing paradigm, which typically assumes that complement proteins primarily target senescent or dysfunctional RBCs during normal homeostasis [17, 19]. This view is based on the interaction of complement proteins with natural autoantibodies that bind to the RBC membrane, contributing to increased C3b formation in aging cells. However, this perspective is focused on the classical pathway [17], overlooking the alternative pathway as a major source of C3b deposition on healthy RBCs throughout their lifespan.

In contrast, our model suggests that the primary role of the complement system is not to facilitate the removal of senescent or dysfunctional cells during normal homeostasis. Instead, it accelerates the clearance of functional RBCs during infections. The continuous formation of C3b on the surface of healthy cells is central to our approach, enabling the selective removal of a fraction of circulating RBC during infections, without disrupting homeostasis under normal conditions.

RBC homeostasis is primarily controlled by EPO in response to the body’s oxygen demands. Our model suggests that the complement system introduces a novel layer of regulation to this homeostatic mechanism. RBCs critically affect blood viscosity [88], and this aspect of RBC homeostasis is essential for ensuring tissue perfusion. We propose that the complement system plays a key immune role, enabling the transient adjustment of RBC numbers to reduce blood viscosity and enhance blood flow to infected tissues during acute-phase responses.

This work sheds light on the intricate interactions between RBCs and the complement system, revealing how these dynamics shape immune responses to infection. Our results provide valuable insights into how the immune system balances its homeostatic and defensive functions. These findings also hold potential clinical relevance, paving the way for novel therapeutic strategies targeting RBC homeostatic disorders, such PNH, inflammation anemia, and other complications that arise during immune responses and inflammation.

## Code availability

Numerical simulations have been performed using Wolfram Mathematica. The code used in these simulations is available at the Notebook Archive (https://notebookarchive.org/2025-05-2sa0yis).

